# Efficient Dual-Negative Selection for Bacterial Genome Editing

**DOI:** 10.1101/2020.03.03.974816

**Authors:** Francesca Romana Cianfanelli, Olivier Cunrath, Dirk Bumann

**Affiliations:** Biozentrum, University of Basel, CH-4056 Basel, Switzerland

## Abstract

We describe a versatile method for chromosomal gene editing based on classical consecutive single-crossovers. The method exploits rapid plasmid construction using Gibson assembly, a convenient *E. coli* donor strain, and efficient dual-negative selection for improved suicide vector resolution. We used this method to generate *in frame* deletions, insertions and point mutations in *Salmonella enterica* with limited hands-on time. Similar strategies allowed efficient gene editing also in *Pseudomonas aeruginosa* and multi-drug-resistant (MDR) *Escherichia coli* clinical isolates.

## Introduction

Genetic engineering is fundamental for molecular analysis of genotype-phenotype relationships, and for determining the function of previously uncharacterized genes [1–3]. Site-specific mutagenesis can be achieved using different methods. Traditionally, marker-free genetic manipulations were obtained using consecutive single-crossovers mediated by endogenous recombinases [4, 5]. A suicide vector is first integrated in the desired location using homologous recombination. Bacteria, in which a subsequent second crossover results in loss of the integrated plasmid, can then be selected using counter-selection markers [6–9]. However, counter-selection is often suboptimal resulting in a need to screen many clones for the desired event. Later, the λ-Red recombineering technology, a phage-based homologous recombination system based on linear DNA transfer and an exogenous recombinase, was introduced [8, 10–13]. Scarless mutations can be obtained when combining this method with a counter-selection marker [14–17]. Currently, λ-Red recombineering is the method of choice for introducing genetic manipulations in *S. enterica* and *E. coli* [18] but it has been difficult to implement in several other bacterial species such as *Pseudomonas aeruginosa*. Recently, CRISPR-Cas has revolutionized eukaryotic genome editing [19–21], but this strategy is more cumbersome for bacteria with limited recombination activities [22–24].

Here, we combined several improvements for establishing a time-efficient versatile method for consecutive single cross-overs in multiple bacterial species. We used rapid Gibson assembly of PCR products [25] to generate suicide vectors with dual negative selection with I-SceI and SacB [26, 27] (Fig 1a), which increased counter-selection efficiency to 100% for nearly all tested deletions, insertions and point mutations. We employed an *E. coli* donor strain that simplifies donor removal after conjugation and avoids common problems with contaminating phages [28]. We used different positive selection markers that enable selection in many bacterial species, including MDR pathogens [29]. Combination of these elements yielded a reliable and fast method for genetic engineering of multiple bacterial species that, in concert with a simplified screening procedure, minimized hands-on time and significantly accelerated genome editing in our lab.

**Figure 1:**
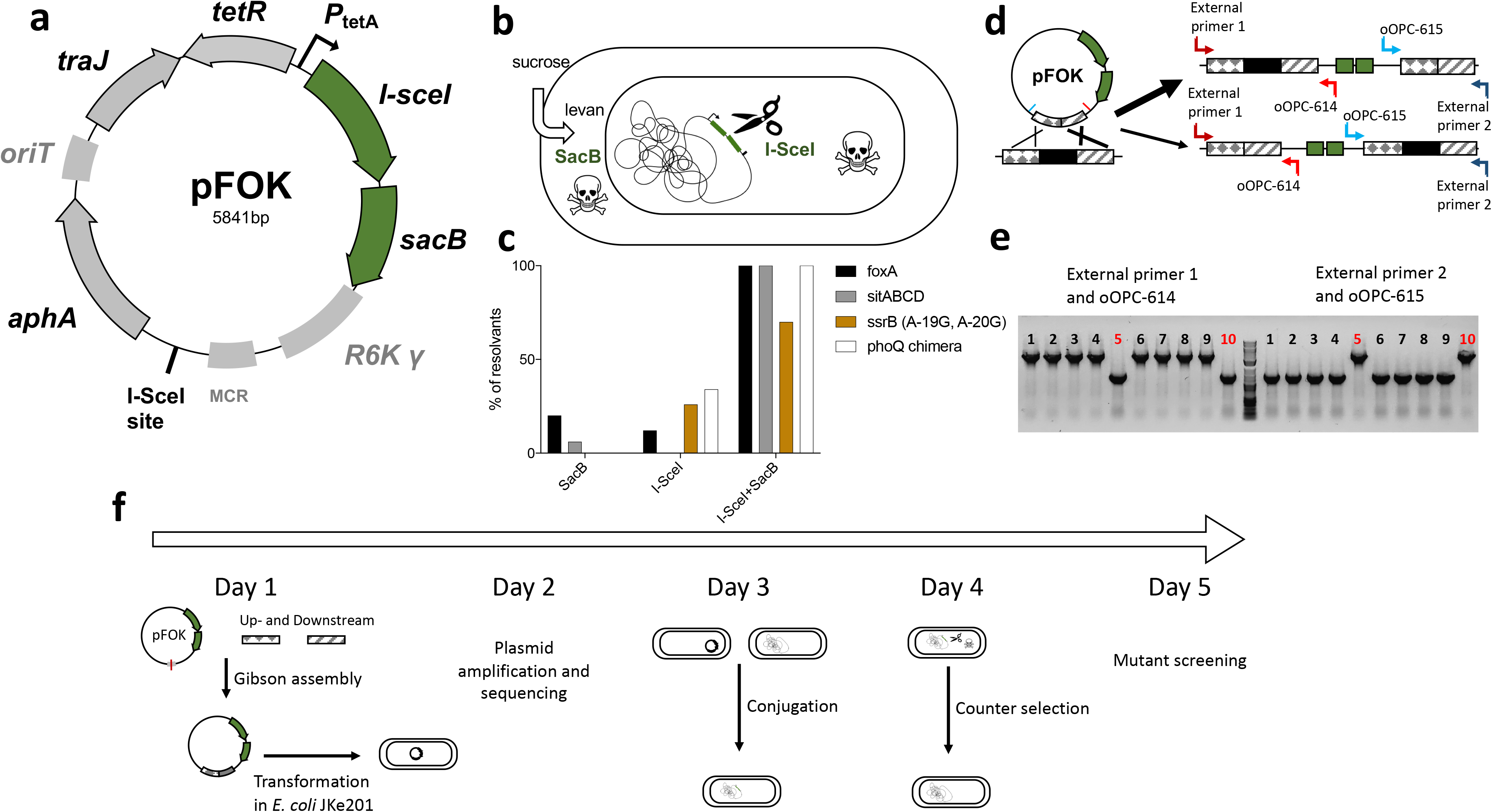
An optimized method for genome editing in *Salmonella enterica*. **a**) Map of suicide plasmid pFOK (*aphA*, aminoglycoside phosphotransferase gene conferring resistance to kanamycin*; I-sceI* gene encoding meganuclease; *oriT*, origin of conjugational transfer; *P*_tetA_, *tetA* promoter, R6K γ *ori*, pi-dependent origin of replication; *sacB*, levansucrase gene; *tetR*, tetracycline repressor gene; *traJ*, transcriptional activator for conjugational transfer genes; MCR, multi cloning region containing <u>at least</u> EcoRI, BamHI, SacI, XhoI and NotI). **b-c**) Negative selection with SacB and I-SceI. **b**) Mechanisms of negative selection for SacB and I-SceI, **c**) Selection efficiency for various chromosomal loci (*foxA* deletion, *sitABCD* deletion, *ssrB* point mutation and *phoQ* chimeric insertion [35]) using either SacB or I-SceI, or a combination of both. **d-e**) Identification of recombination biases favoring one flanking region. **d**) schematic representation of preferential recombination in the right flanking region. External primers (here primer 1 and 2) together with plasmid-specific primers (here primer oOPC-614 and oOPC-615) can be used to screen co-integrant clones to reveal such bias. **e**) Recombination bias for *foxA* gene manipulation. PCR results of ex-conjugant screening using the primer pair 1 and oOPC-614 (left panel here oOPC-396/614) and primer 2 and oOPC-615 (right panel here oOPC-397/615). Rare ex-conjugants (here clone 5 and 10) with recombination in the non-preferred flanking region are used for subsequent counter-selection. **f**) Timeframe with brief summary of daily steps.

## Methods

### Media and strains

Bacterial strains were cultured in lysogeny broth Lennox (LB) (tryptone 10 g/L, yeast extract 5g/L and NaCl 5g/L) medium. *E. coli* JKe201 [28] was cultured in the presence of 100 µM of diamino pimilic acid (DAP) (Sigma Aldrich D1377-10G). *Salmonella enterica* serovar Typhimurium SL1344 was cultured in LB in the presence of 90 µg/ml streptomycin (Sigma-Aldrich S9137-100G*). E. coli* EC01 [29] and *P. aeruginosa* PA14 were cultured in LB. For preparing chemical competent cells, fresh LB medium was inoculated at OD_600nm_ 0.01 with an overnight culture of JKe201 and grown until OD_600nm_ 0.4-0.6. Bacteria were washed twice with 25 ml of ice-cold CaCl2 100 mM (Sigma Aldrich C1016-500G) containing 15% of glycerol (AppliChem, A1123,1000). Bacteria were resuspended in 5 ml ice-cold CaCl_2_ 100 mM containing 15% of glycerol and 200 µl aliquots were frozen and stored at −80°C. Super-Optimal broth with Catabolite repression (SOC) (tryptone 20 g/L, yeast extract 5 g/L, NaCl 0.5 g/L, KCl 0.186 g/L, MgSO_4_ 4.8 g/L and glucose 3.6 g/L) medium was used after heat shock. Kanamycin (Roth T832.4) at a final concentration of 50 µg/ml and gentamicin (Gibco 15750-037) at a final concentration of 15 µg/ml were used to select *E. coli* transformants. For positive selection, kanamycin (Roth T832.4) at a final concentration of 50 µg/ml, gentamicin (Gibco 15750-037) at a final concentration of 30 µg/ml, or potassium tellurite (Sigma P0677) at a final concentration of 10 µg/ml, were used to select for *S. typhimurium, P. aeruginosa* and *E. coli* merodiploids, respectively. For counter-selection plates, LB-no salt (10 g/L tryptone, 5 g/L yeast extract) containing 20% (w/v) sucrose (Sigma-Aldrich 84097-1KG), 15 g/L agar and 0.5 µg/ml anhydrous tetracycline (AHT) (Sigma-Aldrich 37919-100MG-R) were used.

### Generation of the suicide vectors

Primers for generating pOPC-001, pOPC-003 and pFOK are reported in Supplementary Table S1. pOPC-001 was obtained by combining the kanamycin resistance cassette and the I-SceI restriction site from pWRG717 [30], the origin of replication (R6Kγ) and origin of transfer (oriT) from pGP704 [6, 31] and the *tetR* and *I-sceI* locus from pWRG730 [30] using Gibson assembly. pOPC-003 was generated by replacing the *tetR* and *I-sceI* locus from pOPC-001 with *sacB* from pEXG2 [32]. pFOK was generated by inserting *sacB* amplified from pOPC-003 downstream of the *I-sceI* gene on pOPC-001. pFOG was generated by replacing *aphA* of pFOK by *acc(3)-I*. pFOKT was generated by insertion of *tpm* [33] between *aphA* and the multi cloning region (MCR).

### Amplification of the upstream and downstream regions

Flanking primers with a 40 bp overlap were designed to amplify 700bp up- and downstream of the gene of interest using SnapGene® (version 4.0.3) with the Gibson Assembly tool (Supplementary Table S1). Fragments were amplified using a high-fidelity polymerase mix (KOD Hot Start Master Mix, Millipore) and purified on a 1% agarose gel. Vectors were purified from overnight cultures using a plasmid miniprep kit (ZymoPUREtm, ZymoResearch). Vectors were digested using EcoRI-HF and BamHI-HF (New England BioLabs) for 1h at 37°C, or PCR-amplified, and purified on agarose gel. Final concentrations of amplificated fragments and digested vectors were measured using a microvolume spectrometer (Colibri®).

### Gibson assembly and chemical transformation

Plasmids generated in this study are listed in Supplementary Table S2. Gibson assembly reaction was performed as described [25]. The reaction mix contained 50 ng of each up- and downstream fragments and 150 ng of suicide vector, and Gibson assembly mix 1x (New England BioLabs) in a total volume of 10 µl. The reaction mixture was incubated at 50°C for 20 min. 5 µl of the reaction mixture was added to a 100 µl aliquot of *E. coli* JKe201 heat-shock competent bacteria and incubated for 30 min on ice. After a heat shock at 42°C for 30 secs followed by 2 min on ice, bacteria were resuspended in 1 ml prewarmed SOC medium containing 100 µM of DAP and incubated for 1 h at 37°C. Transformants were selected on LB-Lennox agar plates containing 100µM DAP (required by JKe201 for growth) and either 50µg/ml kanamycin, 15µg/ml gentamicin, or 10µg/ml of tellurite. Clones grown on tellurite resulted in black colonies. Clones were screened using PCR with primers oOPC-614 and oOPC-615 (Supplementary Table S1).

### Conjugation and selection of the first homologous recombination event

The recipient *S.* Typhimurium and *E. coli* strains were inoculated in 2 ml of LB containing no antibiotics at 37 °C. *P. aeruginosa* was inoculated in 2 ml LB without antibiotics at 42 °C. The donor *E. coli* strain was inoculated in 2 ml of LB containing 100 µM DAP only without antibiotic. 500 µl each of overnight cultures of the donor *E. coli* strain and the recipient strain were washed with LB, mixed and centrifuged together. The pellet was resuspended in 50 µl LB and deposited on 22 mm filter membranes with 0.45 μm pores (Millipore, Merck) on a pre-dried LB agar plate. After mating for 6 h at 37°C, bacteria were scraped from the filter membrane and resuspended in 1 ml LB. Merodiploid *S.* Typhimurium (pFOK) were selected on LB plates containing 90µg/ml streptomycin and 50µg/ml kanamycin at 37°C for at least 16h. *E. coli* (pFOKT) and *P. aeruginosa* (pFOG) merodiploid were selected on LB plates containing 10µg/ml tellurite or 30µg/ml gentamicin, respectively.

### Counter-selection of the second homologous recombination event

At least three trans-conjugant colonies were combined and grown for 4 h at 37°C in 2 ml of LB. Bacteria were then streaked on freshly prepared LB-no salt agar plates [24] containing 20% sucrose and 0.5 µg/ml AHT. Plates were incubated at 28°C protected from light for at least 24h. Colonies were screened for the desired mutation using PCR with external primers (Supplementary Table S1). Mutants were confirmed by DNA-sequencing (Microsynth.ch).

## Results

Our goal was a rapid and efficient genetic editing method with minimal hands-on time. For this purpose, we combined rapid plasmid construction using Gibson assembly [25], a phage-free, *pir-*carrying (for propagation of *R6Kγ* plasmids), diaminopimelic acid (DAP)-dependent *E. coli* donor strain JKe201 [28] for plasmid amplification and conjugation, with subsequent facile removal of donor in absence of DAP, and an improved dual-negative counter-selection. We generated three suicide vectors from PCR fragments with automatically designed primers using Gibson assembly [25]. Each vector carries commonly used genetic elements for conditional propagation (“suicide vector” with pi-dependent replication from *R6Kγ*), conjugation (*oriT*, *traJ*) and selection for two sequential single-crossovers. For the first positive selection, we developed three different backbones with either; *aphA* conferring resistance to kanamycin (pFOK-Fig. 1a); *aac(3)-I* conferring resistance to gentamicin (pFOG-Fig. 2a); or a plasmid carrying two positive selections *aphA* conferring resistance to kanamycin and *tpm* conferring resistance to tellurite (pFOKT-Fig. 2b) [33].

**Figure 2:**
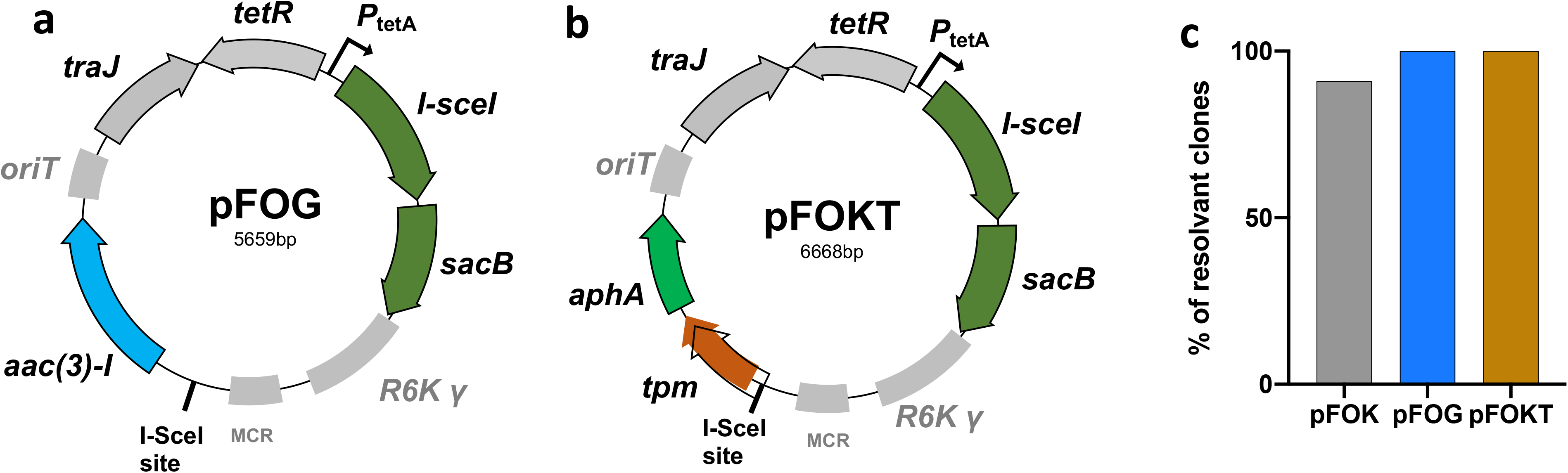
Alternative suicide plasmids. **a**) Map of suicide plasmid pFOG (pFOK analogue except for *aac(3)-I*, conferring resistance to gentamicin) **b**) Map of suicide plasmid pFOKT (pFOK analogue except for *aphA*, conferring resistance to kanamycin <u>and</u> *tpm*, conferring resistance to tellurite) **c**) Selection efficiency for pFOK (*S. enterica*), pFOG (*P. aeruginosa*) and pFOKT (*E. coli*) using the double counter-selection SacB and SceI-I.

A major limitation to efficient genetic editing using two consecutive single-crossovers has been inefficient counter-selection of the second recombination, partially caused by inactivating mutations in the counter-selection marker. We tested counter-selection efficiency in multiple *Salmonella* loci using the commonly used markers *sacB* or *I-sceI*. *sacB* codes for levan sucrase, which confers sensitivity to sucrose because of accumulation of the toxic product levans in the periplasm [26]. *I-sceI* codes for the restriction enzyme I-SceI, which causes lethal DNA double-strand breaks when a I-SceI recognition sequence is present on the genome [27] (Fig. 1b). To assess counter-selection efficiency of SacB or I-SceI singly, we generated plasmid variants (pOPC-001 and pOPC-003) differing just in the counter-selection. Counter-selection was suboptimal for both markers with marker-free clones representing none or only a minority of the recovered colonies (Fig. 1c). Consequently, many colonies had to be tested for finding the desired clones. To overcome this problem, we generated a new suicide vector, pFOK, combining both *sacB* and *I-sceI* under the regulatory control of the TetR regulator. Cells carrying this conditional dual-negative selection cassette showed efficient selection in presence of sucrose and anhydro-tetracycline, yielding a large majority of resolvants that had successfully cured pFOK from their chromosome (Fig. 1c). A similar dual-negative selection has been previously described for Gram-positive bacteria [34].

To expand our gene manipulation method to other bacterial species, including those for which λ-Red recombineering has not yet been established, we used alternative positive selection markers. This included *aac(3)-I*, coding for a aminoglycoside N-acetyltransferase that confers resistance to gentamicin. pFOG in which *aphA* was replaced by *aac(3)-I*, can be used as an alternative to pFOK in bacteria, including *Pseudomonas*, which are resistant to kanamycin but susceptible to gentamicin. We confirmed the utility of pFOG by deleting the *mexAB* operon in *Pseudomonas aeruginosa* (Fig. 2c). As another alternative, we combined *aphA* with a second positive marker, *tpm*, yielding suicide vector pFOKT. *tpm* codes for a thiopurine-S-methyltranferase conferring resistance to tellurite [33]. This plasmid can be used for multi-drug resistant (MDR) bacteria for which the choice of positive selection markers is limited [29]. To limit toxic exposure to volatile dimethyl telluride, we used kanamycin for suicide vector generation and used tellurite only for the positive selection of ex-conjugants. We confirmed the utility of pFOKT by deleting *tolC* with high efficiency in a multi-drug resistant clinical *Escherichia coli* isolate [29] (Fig. 2c).

In some cases, the second single-crossover had a high bias for resolution to wild-type loci (instead of the desired mutant). This was usually due to differences in recombination frequency between the two flanking regions. PCR primers (oOPC614 and oOPC615) that bind in the plasmid, combined with chromosomal primers outside the flanking regions in the merodiploids (Fig. 1d-e), enabled detection of such biases. For these cases, we selected ex-conjugants in which the first single-crossover had occurred in the non-preferred flanking region. In these clones, we often observed frequent resolution to mutant loci during the second single-crossover.

Altogether, the whole protocol from initial plasmid construction to scar-less sequence-verified mutant strains was completed within five working days with minimal hands-on time (Fig. 1f). We have generated more than 30 *Salmonella*, *Pseudomonas* and *Escherichia* gene deletions and point mutations using this method.

## Conclusions

Our plasmids and protocols provide facile time-efficient methods for genetic engineering in multiple bacterial species including MDR clinical isolates. The dual negative selection mitigates the major pitfall of consecutive single-crossovers, the poor selection of resolvants after the second recombination. Gibson assembly enables rapid construction of plasmids with PCR fragments with no need for enzyme digestion and ligation, and no limitations due to restriction sites. As another advantage, our approach relies on endogenous RecA, but not the heterologous, powerful lambda-red recombinase, which might minimize the risk of secondary mutations. Purifying mutated loci by generalized phage transduction may thus not be required. Our method also employs conjugation instead of electroporation (as required for lambda-red methods), which minimizes culture volumes and hands-on time. We anticipate that this method might be broadly applicable to additional bacterial species, including those for which recombineering has been difficult to implement.

## Acknowledgements

We thank all group members for their helpful feedback.

## Author Contributions

D.B. designed the study with input from F.R.C and O.C.; F.R.C. and O.C. constructed plasmids and mutants; F.R.C. and O.C. wrote the manuscript with early input from D.B. and subsequently all authors provided advice and approved the final manuscript.

